# In the AlphaFold era, when is experimental phasing of protein crystals still required?

**DOI:** 10.1101/2024.07.19.604295

**Authors:** Ronan M. Keegan, Adam J. Simpkin, Daniel J. Rigden

## Abstract

The availability of highly accurate protein structure predictions from AlphaFold 2 (AF2) and similar tools has hugely expanded the applicability of Molecular Replacement (MR) for crystal structure solution. Many structures solve routinely using raw models, structures processed to remove unreliable parts or models split into distinct structural units. There is therefore an open question around how many and which cases still require experimental phasing methods such as single-wavelength anomalous diffraction (SAD). Here we address the question using a large set of PDB deposits that were solved by SAD. A large majority (87%) solve using unedited or minimally edited AF2 predictions. A further 17 (4%) yield straightforwardly to MR after splitting of the AF2 prediction using Slice’N’Dice, although different splitting methods succeed on slightly different sets of cases. We also find that further unique targets can be solved by alternative modelling approaches such as ESMFold (four cases), alternative MR approaches such as ARCIMBOLDO and AMPLE (two cases each), and multimeric model building with AlphaFold-Multimer or UniFold (three cases). Ultimately, only 12 cases, or 3% of the SAD-phased set did not yield to any form of MR tested here, offering valuable hints as to the number and characteristics of cases where experimental phasing remains essential for macromolecular structure solution.

## 1. Introduction

In recent times, Molecular Replacement (MR) has become the predominant mode of solution of the phase problem in macromolecular structure determination. MR depends on identifying a search model that is sufficiently similar to at least part of the new crystal structure, allowing its computational placement in the asymmetric unit of the new structure. An initial set of calculated phases can then be determined allowing production of initial electron density maps which may then ultimately allow for structure tracing and refinement. Conventionally, the search model has typically derived from the experimental structure of a quite closely related homologous protein, but the area has also been fertilised by structural bioinformatics. This latter field has encompassed the testing of small substructures identifiable by secondary structure prediction (Rodríguez *et al*., 2009), to diverse approaches for the definition of useful substructures in more distantly related homologues (Sammito *et al*., 2014; Rigden *et al*., 2018) to exploitation of successive waves of *ab initio* structure modelling methods (Bibby *et al*., 2012; Simkovic *et al*., 2016; Simpkin *et al*., 2019).

The latest deep learning-based structured prediction methods, principally AlphaFold 2 (AF2; (Jumper *et al*., 2021)), have had a profound effect on MR. It was immediately recognised at and after the CASP14 structure prediction experiment (Millán *et al*., 2021; Pereira *et al*., 2021; McCoy *et al*., 2022) that their unprecedented accuracy made many targets that would previously have been difficult or intractable, principally those of significantly novel folds, readily soluble by MR. Often an unmodified AF2 model would succeed as a search model, but the importance, in some cases, of removing less confident (often disordered) portions of models and/or splitting multi-domain structure predictions into individual domains, was quickly recognised (Oeffner *et al*., 2022; Simpkin *et al*., 2022). A study of 215 recent PDB deposits that had been solved by single-wavelength anomalous diffraction (SAD) (and which were hence assumed to comprise structures not readily soluble by MR) found that 87% were easily solved using AF2 models, with only seven ultimately failing in MR (Terwilliger *et al*., 2023). Such results have encouraged the integration of AF2 into structure solution pipelines (Krissinel *et al*., 2022; Poon *et al*., 2024; Simpkin, Caballero *et al*., 2023).

Here we address to what extent search models from AF2 and other approaches can solve recent PDB (Berman *et al*., 2003) deposits that had been solved by single-wavelength anomalous diffraction (SAD). The set of 408 cases, deposited between the 27th of July 2022 and the 4th of October 2023, is much larger than tested in previous work but, it should be remembered, represents only a minority of PDB accessions: in the same period 10,084 deposited structures list that they were determined using MR. And crucially, as well as testing the performance of different AF2 variants (the original DeepMind version (Jumper *et al*., 2021) and ColabFold (Mirdita *et al*., 2022)), we quantify the importance of (automated) domain splitting, as well as illustrating the key value in difficult cases both of other software (such as ESMFold (Lin *et al*., 2023) and ARCIMBOLDO (Rodríguez *et al*., 2009); and of alternative approaches (eg multimeric search model building). All in all, just 3% of the SAD test set were not solved, providing a new picture of the size and characteristics of recent PDB deposits for which experimental phasing was likely essential for structure solution. We find that targets with few homologues in the database are challenging, as expected, and that proteins with predominantly alpha secondary structure, especially coiled-coils, are also over- represented in difficult cases. While experimental phasing is widely used for other purposes (El Omari *et al*., 2023) e.g. localisation of ions, these results help define the future targets which should be prioritised for this typically more expensive and laborious mode of structure solution.

## 2. Methods

### 2.1 Selection of target sequences

The targets selected were PDB structures that had been solved by single-wavelength anomalous diffraction (SAD) and deposited between the 27th of July 2022 and the 4th of October 2023. This yielded 408 cases, two of which (8ec9 and 8cuf) contained peptides with multiple modified or unnatural amino-acids and so were excluded (see Supplementary Material for a list of PDB IDs).

### 2.2 Modelling and characterisation of target sequences

Target sequences were modelled using DeepMind AlphaFold 2 (here called AF2;(Jumper *et al*., 2021)) and ColabFold (CF; (Mirdita *et al*., 2022)). These methods employ the same trained network but differ in other aspects, principally in the mode of generation of Multiple Sequence Alignment (MSA), a key input. DeepMind AF2 uses HHblits (Remmert *et al*., 2011) and jackhmmer (Johnson *et al*., 2010) to search the Big Fantastic Database (BFD), Uniclust30, Uniref90 and MGnify databases while ColabFold uses the mmseqs2 application programming interface (Steinegger & Söding, 2017) allowing rapid search of metagenomic databases. By default, AF2 used structural templates (homologous structures identified in the PDB), whereas CF did not.

Alignment depth was expressed as Neff, a measure of effective number of sequences for MSAs (Wu *et al*., 2020; Jumper *et al*., 2021) using an 80% sequence identity threshold and normalised by target length, hence Neff/length. Targets were classified using the secondary structure information in the deposited PDB structures, defining them as all-α (100% of regular secondary structure was α-helix), mostly α (>65% α), mixed, mostly β (>65%), or all- β (100% of regular secondary structure was β-sheet). Socket2 (Kumar & Woolfson, 2021) was used to evaluate if targets contained coiled-coil domains within them, as it is noted that AF2 struggles to accurately predict these domains (van Breugel *et al*., 2022).

### 2.3 Processing of structure predictions into search models

AF2 predictions are accompanied by residue level confidence estimates expressed as predicted local difference distance test (pLDDT) numbers on a scale of 0-100 where values of 70 or higher are considered as moderate-to-high confidence (Jumper *et al*., 2021). The AF2 and CF models were trialled unmodified or after trimming to remove residues with a pLDDT confidence estimate of less than 70. In each case, the pLDDT values of the structure were converted to pseudo B-factors (Oeffner *et al*., 2022). For some multi-domain targets where these trials failed, Slice’N’Dice (SnD; (Simpkin *et al*., 2022)) was used to split the model into distinct structural units. A description of how this was done is given in the section 2.5.2 below (MR and refinement). SnD allows for the selection from a number of algorithms for structural domain splitting: here, the Birch algorithm was used by default (balanced iterative reducing and clustering using hierarchies; (Zhang *et al*., 1996)), with splitting based on the AF2 Predicted Aligned Error (PAE; (Jumper *et al*., 2021)) tested in some cases. The Birch algorithm amounts to applying clustering on the C-alpha coordinates of the input structure whereas the PAE-based method generates clusters from the PAE matrix output from AF2. The PAE matrix displays the predicted alignment error to the true structure for each residue pair. Contiguous regions of this matrix having a low PAE usually represent a domain or structural unit. Where single molecule AF2 and CF predictions failed to yield a solution, ESMFold models (Lin *et al*., 2023) were generated and subjected to the same pLDDT-based trimming and splitting if necessary. Multimeric search models were also generated for these cases using AlphaFold-Multimer (Evans *et al*., 2022) or Unifold- symmetry (Li *et al*., 2022) and subjected to the same pLDDT-based processing.

### 2.4 Secondary structure-based search models

ARCIMBOLDO_LITE (Rodríguez *et al*., 2009) was used to try to solve selected α-helix rich targets. AMPLE helical ensemble search models (Sánchez Rodríguez *et al*., 2020) were also trialled in some cases.

### 2.5 Molecular Replacement and refinement

#### 2.5.1 First pass: Automated tests

Predictions were created using local installations of both AF2 (version 2.3.2) and CF (version 1.5.1) and passed into an automated structure solution pipeline. This pipeline was based on applications from the CCP4 Suite (version 8.0.016, (Agirre *et al*., 2023)). Models were prepared as search models for MR using SnD to truncate residues with a pLDDT<70 but not performing any splitting. MR was carried out using Phaser (McCoy *et al*., 2007; Read & McCoy, 2016). The resulting solution was subjected to 100 cycles of jelly-body refinement in Refmacat (Yamashita *et al*., 2023). Phaser was run with default options. The assumed similarity of the search models to the target was set to 1.2 Å r.m.s.d. Assessment of the solution was done using a map correlation coefficient (map CC) calculated between the placed model(s) and a map calculated from the deposited structure. This was performed using phenix.get_cc_mtz_pdb from the Phenix suite (Liebschner *et al*., 2019). A global map CC of greater than 0.25 was considered to be indicative of correct placement for the search model(s). Where a manually-driven structure solution was required (described below), Phaser’s LLG (log likelihood gain) and TFZ (translation function Z-score) were also used to assess the MR solutions. An LLG of better than 64 and a TFZ better than 8.0 for each search model in MR are considered to be indicative of correct placement (Oeffner *et al*., 2018).

#### 2.5.2 Second pass: Manual structure solution

Where the automated tests failed to give a solution meeting the map CC threshold for a solution, test case data and predictions were imported into a CCP4 Cloud project (Krissinel *et al*., 2022) where search model preparation, MR and subsequent steps could be carried out interactively through the interface. If Multimer and ESMFold predictions were tested in these cases, the predictions were generated online using the Deepmind AF2 google Colab server, the Unifold google Colab server and the ESMFold prediction service provided through the ESMAtlas website. These were then imported and prepared for MR in the CCP4 Cloud project. Solution attempts using ARCIMBOLDO were also run through the CCP4 Cloud interface. AMPLE tests were run from the command line. Where model splitting was required, several splits were tested. This typically involved splitting the initial prediction into two, three or more parts with SnD and passing all parts as separate search models to a Phaser MR run. The tree-like project structure layout of CCP4 Cloud made it easy to explore each of the split strategies. In some cases, the parameterisation of Phaser was adjusted to aid the determination of a correct solution. For example, sometimes the assumed r.m.s.d. of the search model to the target needed to be decreased from the chosen default of 1.2 Å to enable the correct placement of search models that were more accurate, but constituted only a small part of the overall scattering content in the crystal. In other cases, only partial solutions could be found in MR, so additional refinement in Refmacat and model building using Modelcraft (Bond & Cowtan, 2022) were run to verify the solution and bring it to a more complete state.

## 3. Results and Discussion

### 3.1 Most recent SAD-phased structures can be solved with unedited or pLDDT-edited AF2 or CF models

We first attempted structure solution using DeepMind AF2 models, either unmodified or after trimming to remove residues with a pLDDT confidence estimate of less than 70. In each case, the pLDDT values of the structure were converted to B-factors as in Methods. Using a map CC of at least 0.25 (see Methods) as a threshold, a large majority of the test cases achieved this criterion- 349/406 = 86 % - solving straightforwardly (Figure 1). Recalling that these are not PDB deposits as a whole, rather those for which SAD was used for phasing, this is a remarkable result that echoes previous work demonstrating the profound impact of AF2 on crystal structure solution by MR (Terwilliger *et al*., 2023).

**Figure 1.**
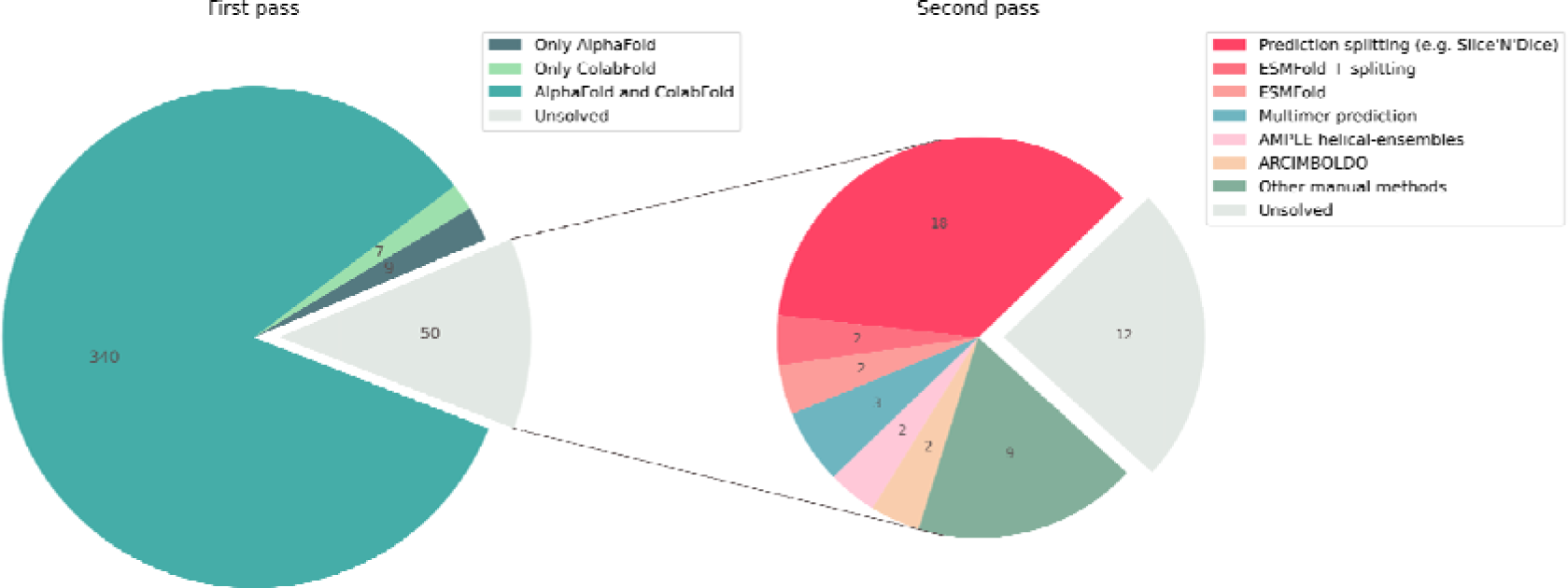
Pie charts showing the results from testing our SAD dataset (406 structures). The pie chart on the left shows the results from the first pass where models from ColabFold and AlphaFold were run in MR with minimal modification (pLDDT ≥ 70 and conversion pLDDT to predicted B-factor conversion), 356 of the 406 structures (87.7%) solved in this first pass. The pie chart on the right shows the results from the second pass where various MR strategies were used on 50 cases that we had been unable to solve in the first pass. A further 38 cases were solved bringing the total to 394/406 (97%).

#### 3.1.1 ColabFold solves a slightly different set to DeepMind AlphaFold 2

The development of the ColabFold implementation of AF2 (Mirdita *et al*., 2022) has contributed significantly to the democratisation of the technology. By replacing the original MSA generation step of AF2 with a quicker approach based on mmseqs2 (Steinegger & Söding, 2017) and, in particular, by provision of a fast API for the MSA generation step, the CF developers have facilitated AF2 modelling on more modest hardware and, notably, on Colab pages. In most cases, AF2 and CF each produce excellent models which can be used largely interchangeably for downstream applications. However, we note here that the different MSAs generated by AF2 and CF can lead to subtly different structures with, in some cases, a crucial impact on MR success (Suppl Table 1).

#### 3.1.2 Inclusion of templates is occasionally required for success

The AF2 pipeline employed here used templates where available. Although not required in most cases, where the available MSA depth and power of the modelling network combine to produce excellent models, AF2 and CF can optionally use information from homologous templates. A dramatic difference in the quality of the AF2 and CF models for 7snr was noted (Figure 2). It could be explained by the exceedingly shallow MSA produced by the CF pipeline (Neff/length = 0.01) which allows for the extraction of only weak evolutionary covariance information. The AF2 model was far superior since, unlike the CF prediction, it used information from the PDB structures (5b0u, 4azn, 5ur9, 4lkp) as templates.

**Figure. 2.**
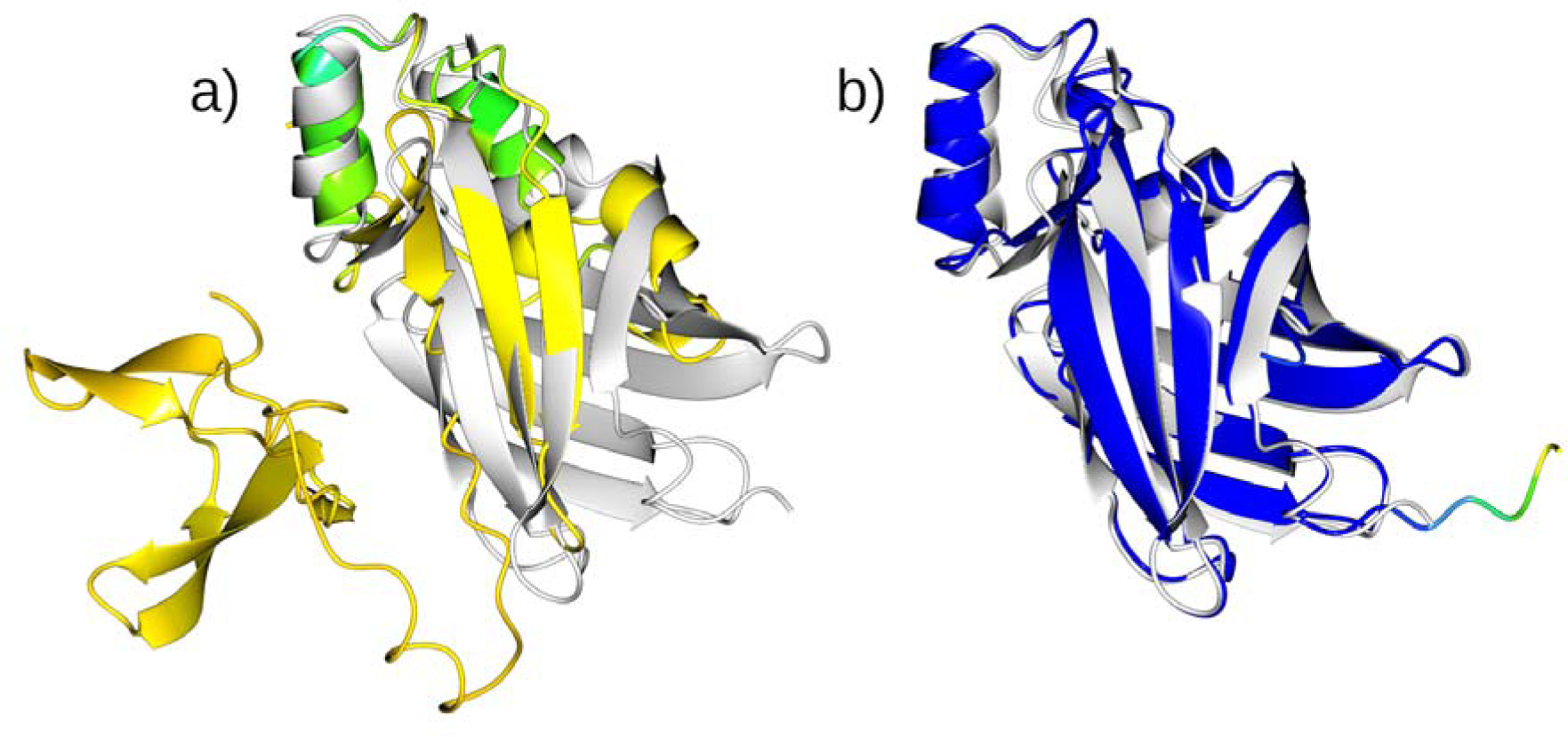
The alignment of the Colabfold (a) and AF2 (b) predictions coloured by pLDDT to the deposited structure for 7snr (2Å, 2 copies in the asymmetric unit, spacegroup P21212) (white). The AF2 prediction benefitted from the use of a template during the prediction process to give a much higher confidence and subsequently more accurate prediction suitable as a search model in MR. All structure figures were created using the CCP4mg application (McNicholas *et al*., 2011).

### 3.2 Most of the remaining 50 structures yield to structure solution using computational methods

The 50 structures not straightforwardly solved above were examined in more detail (Suppl Tables 2-7). As described in 2.5.2, alternative computational approaches were trialled, principally the splitting of AF2 models more finely into larger numbers of structural units. On a case-by-case basis, other approaches included AF2 Multimer, UniFold (especially the symmetry-aware version) and ESMFold. MR with ideal helices in ARCIMBOLDO_LITE (Rodríguez *et al*., 2009) or helical ensembles in AMPLE (Sánchez Rodríguez *et al*., 2020) was also attempted in selected cases. Suppl Tables 2-7 summarise key characteristics of these cases and indicate whether and how successful MR was achieved. Below, we consider some of the main findings and provide illustrative examples in each case.

#### 3.2.1 Automated splitting into two or more units solves many more cases

It is well understood that multi-domain proteins represent a challenge, even for the latest generation of modelling software. Inter-domain information from evolutionary covariance will likely be weaker than intra-domain information and, of course, flexible linkers between domains can lead to proteins sampling conformations with a variety of inter-domain orientations: in such cases the prediction may be valid as biologically accessible, but only a representative conformation that may differ significantly from the crystallised conformation. Recognising the issue, it is relatively commonplace to split domains of larger proteins and the PAE, provided by many methods, provides a readout of confidence in inter-domain orientations that can be used for splitting. Targets that yielded to this approach are listed in Suppl Table 2. Here, we first applied the default PAE-independent Birch algorithm for domain splitting by SnD: this method allows for processing experimental structures and structure predictions from any method.

An example of successful structure solution after Birch domain splitting is shown in Figure 3 for the test case 8ewh. After eliminating low-confidence residues (see Methods), the structure prediction is split into four units which superimpose on the respective domains of the native structure much better than the original prediction. An MR search for the two copies of the overall molecule i.e. the attempted placement of 2 copies of each of the 4 search models created by the splitting, resulted in the correct placement for all parts with an LLG of 3458. Subsequent jelly-body refinement with Refmacat improved the solution to an R/Rfree of 0.29/0.33.

**Figure 3.**
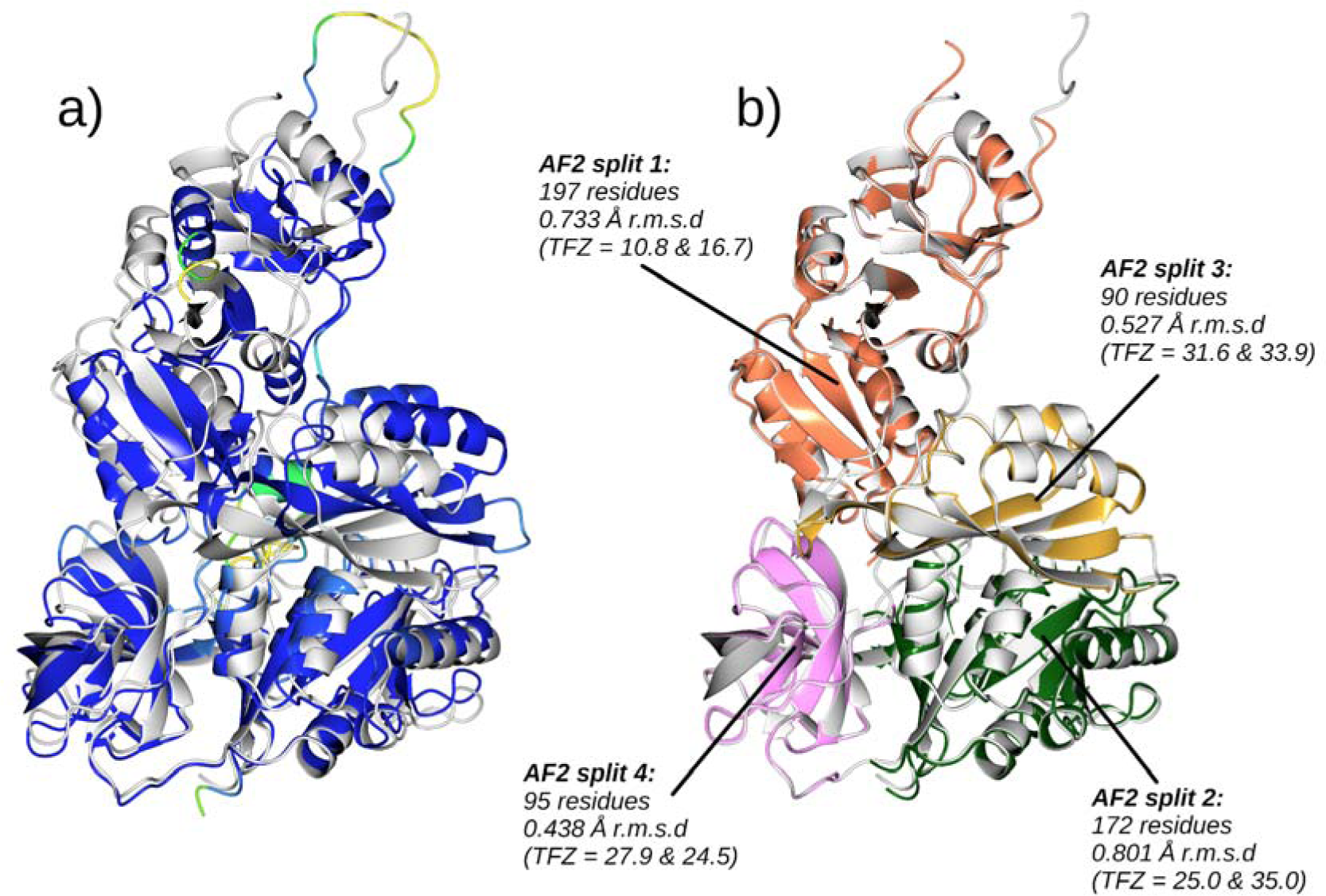
- 8ewh (2.33 Å, 2 copies in the asymmetric unit, spacegroup P21). a) shows the unmodified AF2 prediction (coloured by pLDDT) aligned to the crystal structure (white) using Gesamt to perform the alignment (2.5 Å r.m.s.d.). MR using the full predicted model was not successful. b) shows the alignment to the true structure of a 4-way split of the AF2 prediction after it has been truncated to residues with pLDDT >= 70. The Birch clustering method employed in SnD has been used to cluster the c-alpha atoms into the 4 clusters. Phaser successfully placed all 8 fragments making up the 2 copies of the molecule in the asymmetric unit with a final LLG of 3458. TFZ values for each placed fragment are shown. To illustrate their accuracy, the r.m.s.d. for each split fragment to the crystal structure is also shown. The solution was then refined using 100 cycles of jelly-body refinement in Refmacat giving an R/Rfree of 0.29/0.33. The solution was further improved to an R/Rfree of 0.24/0.30 by automated model building using Modelcraft. This gave a global map CC of 0.772 when compared to the map generated from the deposited data.

#### 3.2.2 The splitting algorithm can be important

Domain partition of protein structures is an active area of research and a variety of approaches and programs are available eg (Wells *et al*., 2024; Kandathil *et al*., 2024; Cretin *et al*., 2022). The default Birch algorithm in SnD has the advantages of being quick and PAE-independent, meaning that it can be applied to any input structure. However, in some cases, use of alternative algorithms is required for success. For example, the target 8bfi was not solved by SnD using Birch-defined partitions, but was solved using PAE-based domain splitting (Figure 4)

**Figure 4.**
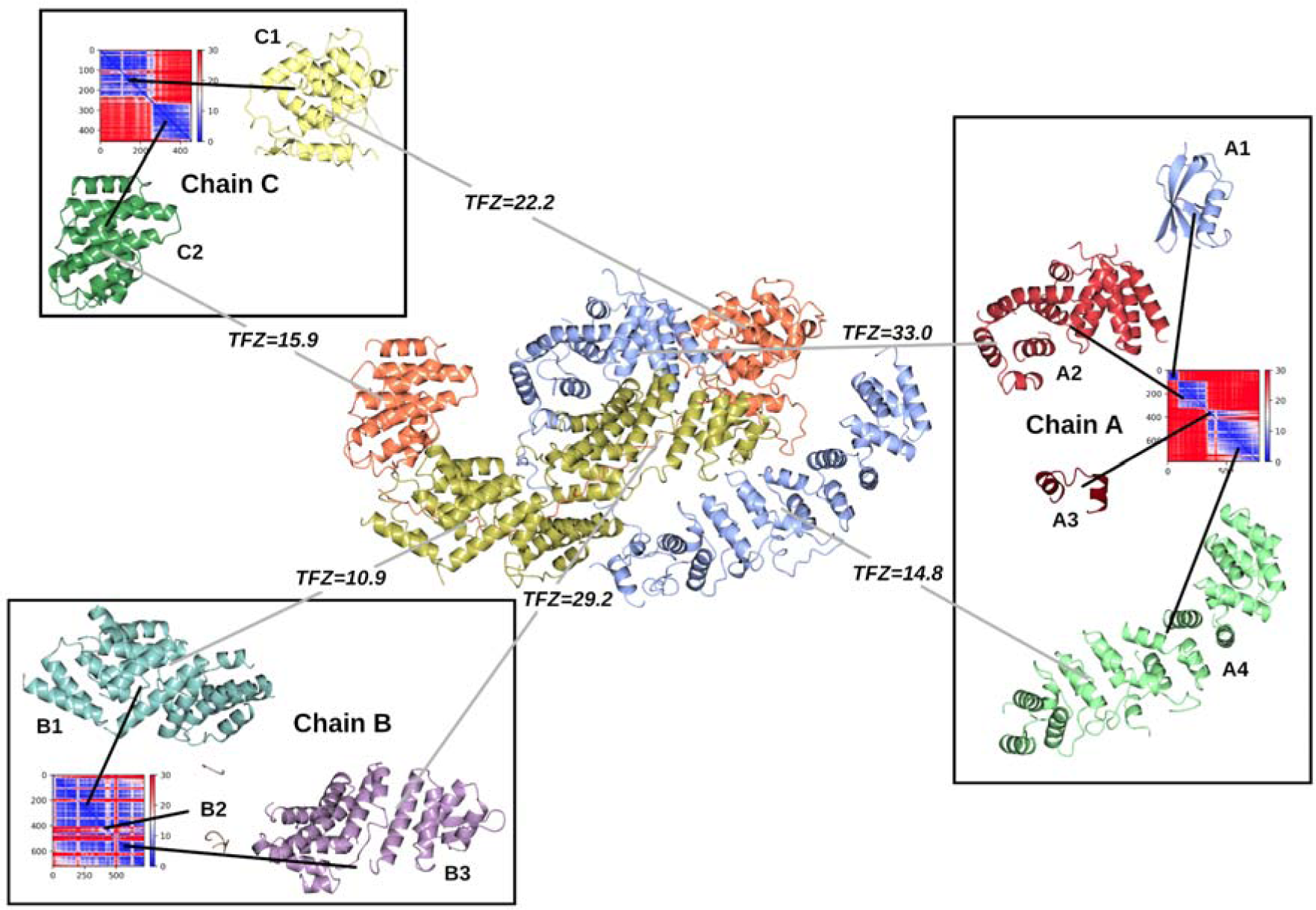
Solution to target structure 8bfi (3 Å, 1 copies in the asymmetric unit, spacegroup P21) using PAE-based domain splitting of Colabfold predictions of each of the 3 chains to create suitable search models for MR. The target structure is shown in the centre, coloured by its 3 chains A (blue), B (gold) and C (orange). Chains A and C have domain-swapped parts leading to large differences in the relative orientation of each of the predicted chain’s domains when compared to the crystallised form. However, utilising the PAE matrices produced by Colabfold (shown for each chain, blue=low error, red=high), the domains can be separated out from the chain predictions to produce accurate search models (shown in the boxes) that can be placed correctly in MR by Phaser . All predicted domains were placed correctly apart from A1, which was not present in the crystal, and A3 and B2 which both constituted a very small fraction of the overall scattering content. The TFZ from Phaser for each search model placement is shown (values above 8 are indicative of correct placement). The final LLG for all of the placed components was 2048.

#### 3.2.3 Alternative model building software can succeed

Recent CASP data have confirmed that AF2 predictions continue to underpin all the most successful structure prediction protocols. However, there is also strong interest in the alternative approach of structure prediction using protein Language Models (pLMs), especially for its much superior predictive speed, and CASP15 indeed showed that in a handful of cases, the best-performing pLM method at the competition produced the most accurate model. Targets that yielded to this approach are listed in Suppl Table 3.

For the case 8bjw, neither the AF2 or CF models succeeded with the approaches we tried, and nor did the target yield to MR with ideal helices in ARCIMBOLDO_LITE. However, a prediction from ESMFold, processed to remove low confidence residues did succeed (Figure 5). In this case, removal of pLDDT<70 residues from the AF2 model left only 16 residues which, given the six copies in the asymmetric unit and moderate resolution, failed in MR. Interestingly, the processed ESMfold result (60 residues, 0.503 Å, r.m.s.d.) is only a little larger and more accurate than the CF-derived search model (45 residues, 1.215 Å r.m.s.d.) but the marginal difference is evidently crucial. The final LLG for the six placed copies of the ESMFold-based search model was 655 with TFZ values for the placed copies ranging from 8.6 to 16.7. With refinement and model building the R/Rfree was 0.22/0.32.

**Figure 5.**
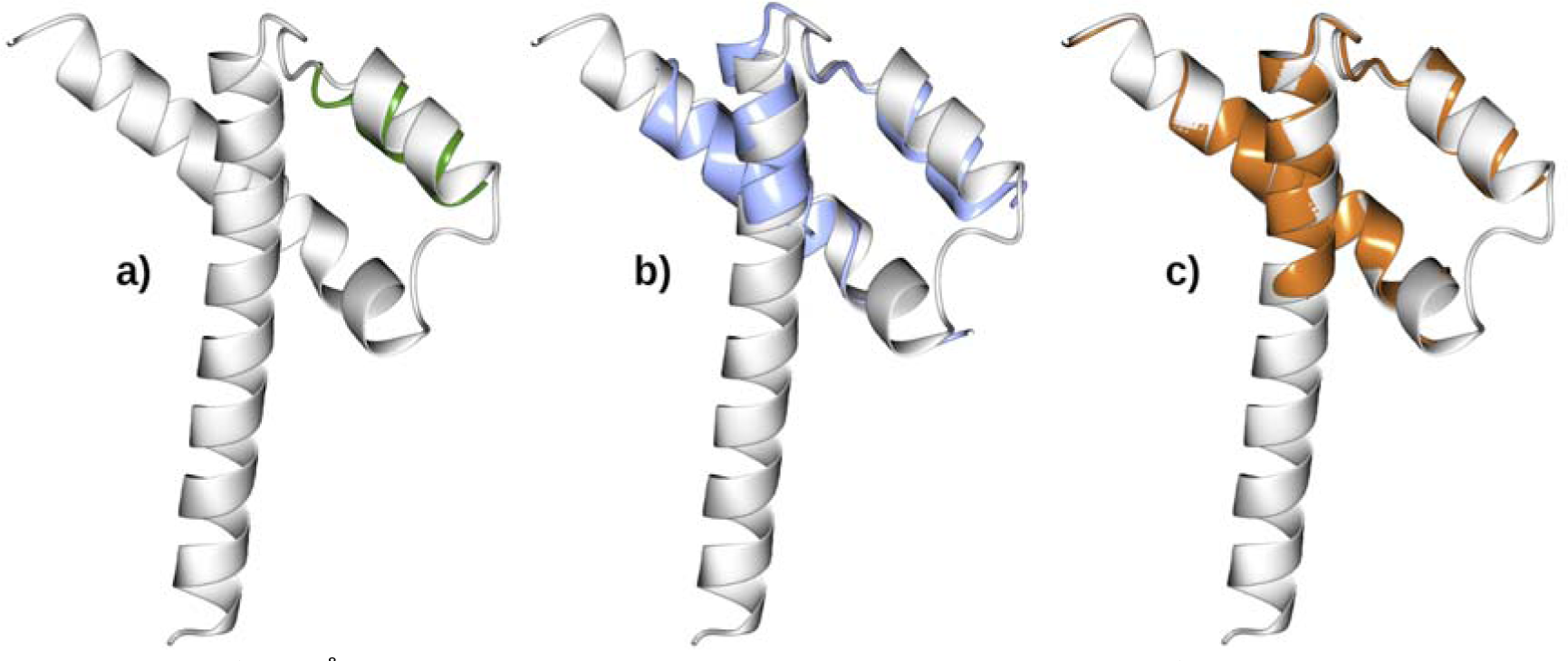
8bjw (2.39 Å, 6 copies in the asymmetric unit, spacegroup P3221). Following model truncation to residues with pLDDT>=70, target structure (white) aligned to a) AF2 prediction (green, 16 residues, r.m.s.d. 1.07 Å), b) Colabfold prediction (blue, 45 residues, r.m.s.d. 1.215 Å) and ESMFold prediction (orange, 60 residues, r.m.s.d. 0.503 Å)

The case of 7qrr is even more remarkable. The AF2 and CF models are each very poor (Figure 6), even when modelling the native trimer (data not shown), presumably due to the low Neff/length values of their input MSAs (0.01 in each case). The ESMFold model, however, is excellent and a remarkable 12 copies can be placed to solve the structure (LLG=2064). Interestingly, 7qrr and 8bjw are both virus proteins whose rapid sequence divergence during evolution is known to make collection of homologous sequences more difficult (Karlin, 2024). Conceivably, in these cases the ESMFold language model training more successfully inferred relationships across these divergent families than were discovered during the MSA generation of AF2 and CF modelling.

**Figure 6.**
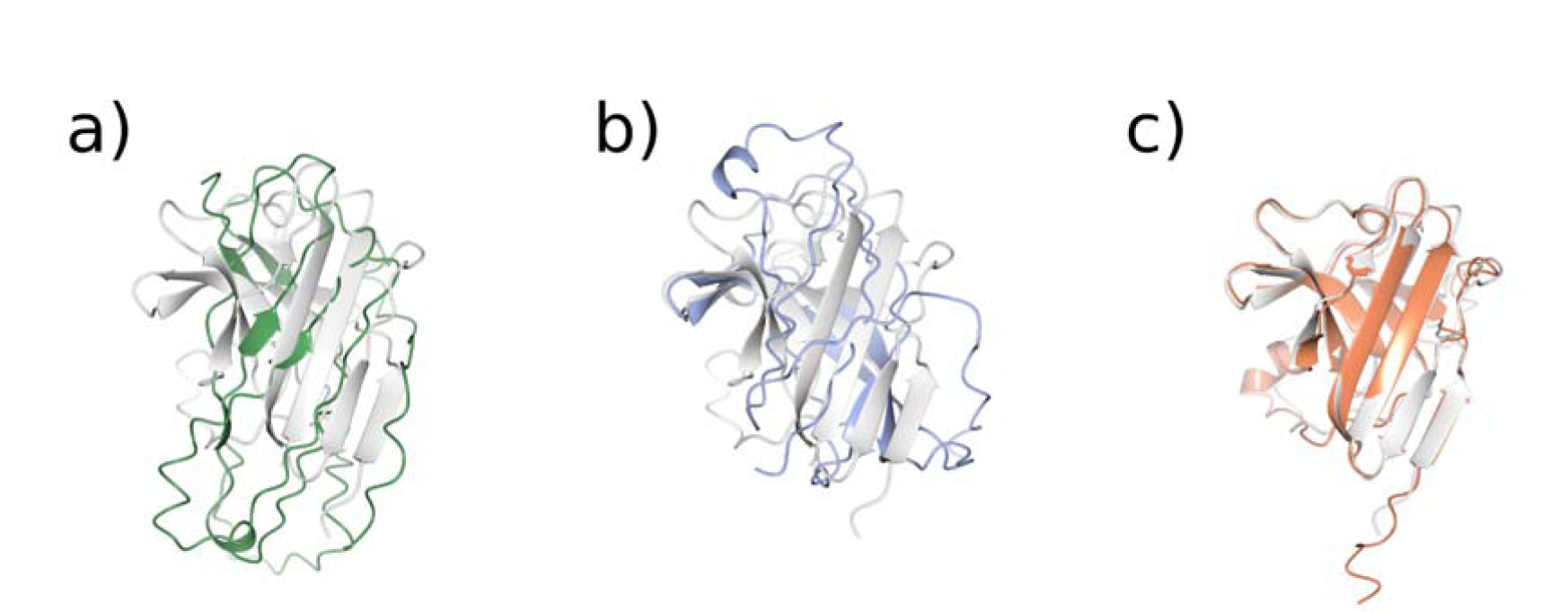
7qrr (1.9 Å, 12 copies in asymmetric unit, spacegroup C121). Target structure (white) aligned to a) AF2 prediction (green, r.m.s.d. 4.123 Å), b) Colabfold prediction (blue, r.m.s.d. 3.636 Å) and ESMFold prediction (orange, r.m.s.d. 1.384 Å).

In the remaining two cases, 8bvl and 8gxl, splitting of ESMfold predictions with SnD was required for structure solution.

#### 3.2.4 Modelling an oligomer can be helpful

In some cases it can be critical that the search model contains a significant proportion of the scattering matter e.g. where data only extends to a relatively low resolution. Thus, construction of oligomeric search models can help in these scenarios, giving the search model a larger signal in the MR search and enabling correct placement to be found more readily. The latest generation of structure prediction software can straightforwardly model protein complexes (Evans *et al*., 2022) and, in some cases (Li *et al*., 2022), can exploit non-crystallographic symmetry that might be apparent, for example, from a self-rotation function.

In the set here, there were three cases where construction of an oligomer was absolutely essential (Suppl Table 4). For example, Figure 7 shows the case 8h8h where a monomeric structure was insufficiently accurately captured. In this case modelling the dimer produced a more accurate model, with the angles between the two helices closer to the native structure. The resulting dimer can be used to solve the structure but required some manual assistance to find all 4 copies. 3 copies were initially placed by Phaser achieving an LLG of 220.

**Figure 7.**
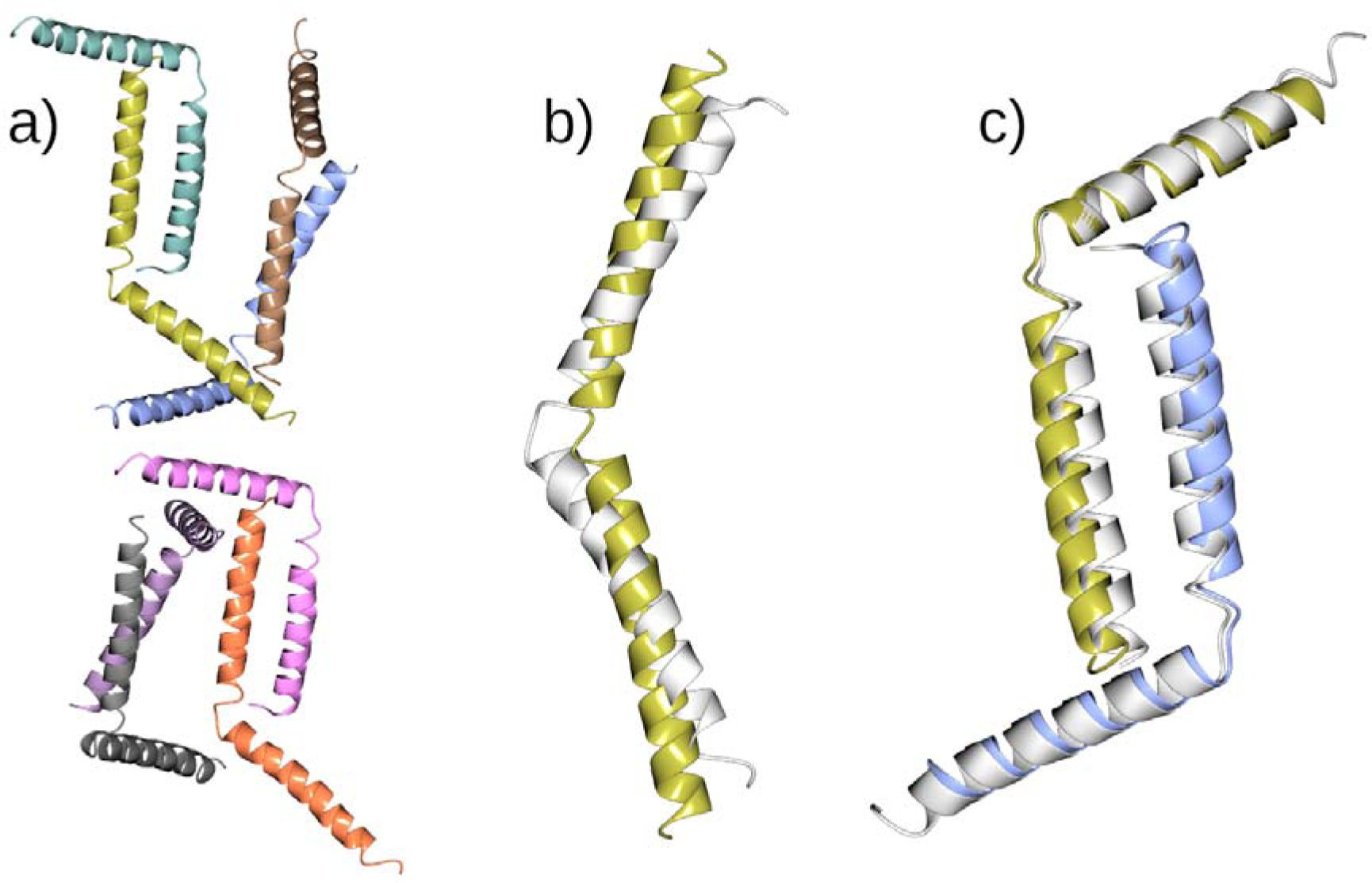
8h8h (2.7 Å, 8 copies in the asymmetric unit, spacegroup P212121). a) shows the entire content of the asymmetric unit, b) is the structural alignment of the AF2 prediction and a single chain from the deposited structure, c) is a structural alignment between a dimer from the deposited structure (white) and a dimeric model generated using Unifold. Both the AF2 prediction and the Unifold dimer have been truncated to residues with pLDDT >= 70. The AF2 prediction failed to yield a solution in MR, either as the whole monomer or as a set of derived search models through splitting at the hinge between the helices. Both the dimer and the monomer produced as part of the Unifold dimer prediction can be used to determine the structure.

Refinement of this solution and subsequent use of one of the refined dimers as a search model, enabled the 4th copy to be placed correctly (TFZ=34.9). The better formed monomer produced in the dimer prediction gives a solution more easily, with all 8 copies found by Phaser, achieving an LLG=1427.

The case of 7rt7, where the crystal contains 6 copies of a 2-chain heterogeneous complex, demonstrates the advantage of using a complex prediction rather than the individual monomers in MR. Although the monomeric search models can be used in MR successfully, the stronger signal for the complex search model containing the 2 chains solves the structure much more quickly (15 minutes vs 2 hours, Figure 8).

**Figure 8.**
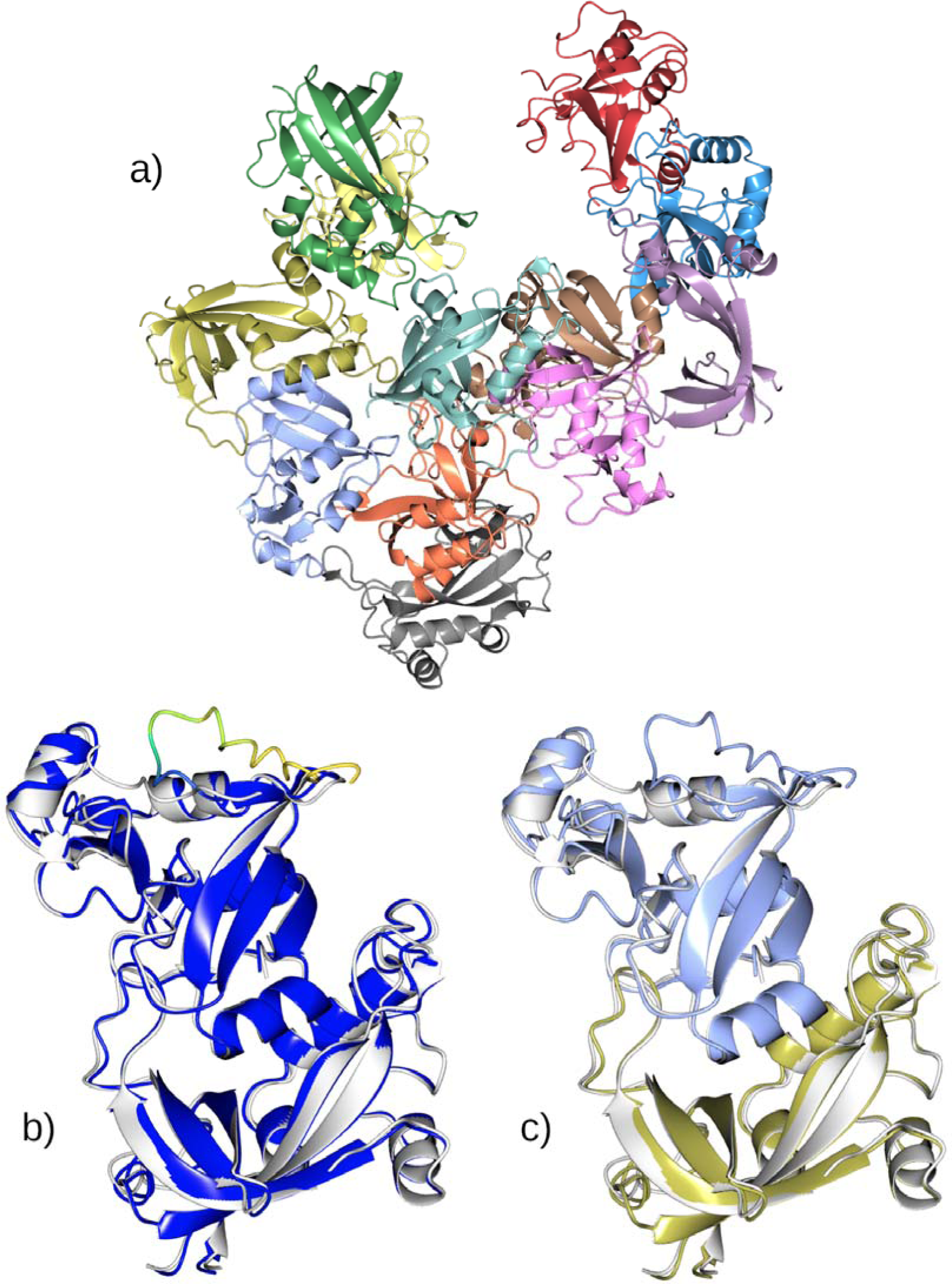
7rt7 (2.29 Å, 12 copies in the asymmetric unit (6+6), spacegroup P3221). The asymmetric unit consists of 6 copies of a heterogeneous complex of 2 chains (a). The AF2 prediction for the complex coloured by pLDDT aligned to the true structure of the complex (white) is shown in b), c) displays the same alignment with the predicted complex coloured by chain. The r.m.s.d. for the aligned complex to target is 0.735 Å. A solution could be found with Phaser using the individual chains as search models (taking 2 hours on 4 cpu cores). However, using the complex search model gave a clearer signal for the correct positioning of the models, enabling a solution to be found in less than 15 minutes.

#### 3.2.5 Secondary structure-based search models are still useful for coiled-coil and other helix-rich cases

As was evident at CASP15 (Simpkin, Mesdaghi *et al*., 2023), even the latest modelling software can struggle with certain cases, most notably those for which few homologues are available by database search (and for which, of course, there is no template in the PDB).

With these proteins, AF2 and similar tools have neither the template nor the evolutionary covariance information required to generate an initial model. Here, it must be recalled that AF2 is not the only tool in the box, and previous generations of unconventional MR software can still be deployed.

We first used ARCIMBOLDO_LITE (Sammito *et al*., 2013) which solved 7twd and 7qwe (Supp Table 5). In the case of 7twd, where each chain is a single, kinked α-helix, the helical twist is incorrectly modelled by AF2 and CF. 7qwe is the structure of a *de novo* protein containing a homo-pentamer. AF2, CF and UniFold models of the single chain or oligomers failed but ARCIMBOLDO_LITE solved the structure (Figure 9).

**Figure 9.**
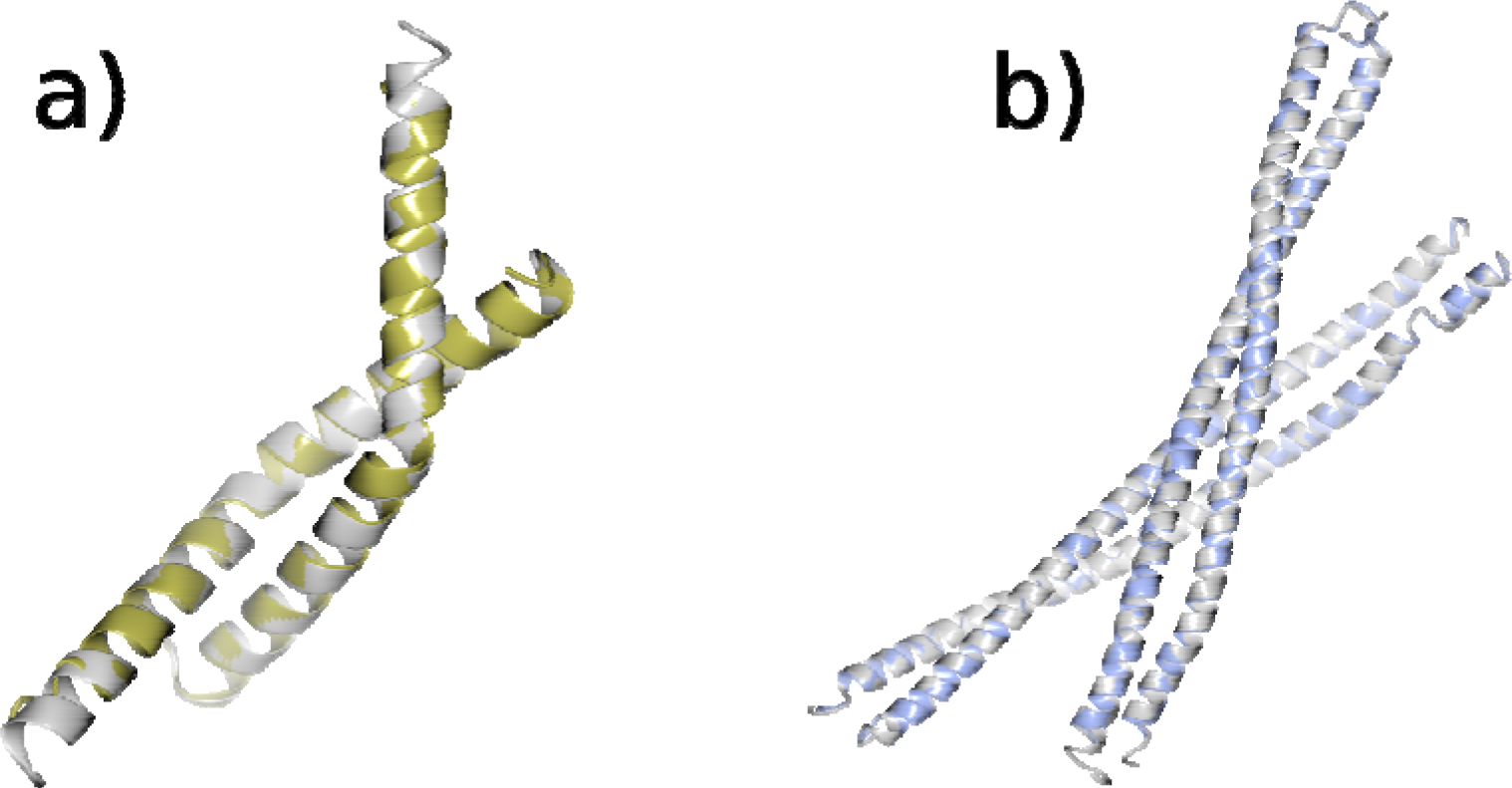
a) The ARCIMBOLDO solution (gold) shown against the crystal structure (grey) for 7twd (2.11Å, 2 copies in the asymmetric unit, spacegroup P6122) b) The AMPLE solution (blue) shown against the crystal structure (grey) for 8fby (2.4Å, 4 copies in the asymmetric unit, spacegroup P212121).

For a handful of cases of structures that were rich in α-helical content but which were not solved any other way, one containing a coiled-coil, we attempted MR with AMPLE using libraries of helical ensembles which have been shown to outperform single ideal helical structures (Sánchez Rodríguez *et al*., 2020). For two targets, 8bc5 and 8fby, successful MR results were thereby obtained with search models of 30 and 25 residues respectively (Supp Table 6). Interestingly, 8bc5 featured as target T1122 at CASP15 where the extreme low solvent content of the crystal was noted as a feature that aggravated the high degree of difficulty imparted by the shallow MSA (Simpkin, Mesdaghi *et al*., 2023).

#### 3.2.6 Further investigation and intervention was required in some cases

Where the straightforward manual processing approach in CCP4 Cloud failed to give a solution or only a partial solution, a more detailed examination of the case was sometimes required (Supp Table 7). One such case is 8u12 (Crystal Structure of Antitoxin Protein Rv0298), which consists of 8 copies of the target protein in the asymmetric unit (Figure 10 (a)) in H3 with a resolution 1.6 Å. 25 residues from the N-terminal of the expected sequence do not appear to have been crystalised, making MR using a predicted model based on this sequence difficult due to packing clashes between residues that are not present. In the initial MR trial, using the entire predicted model from AF2 (Figure 10 (b)), Phaser struggled to find the correct placement for the models due to these packing clashes. This was followed by using SnD to split the model in two (Figure 10 (d)) and attempt to place 8 copies of each part. Using these parts as search models, Phaser gave a more encouraging solution with an LLG of 2528. Closer examination of the result indicated that it had placed 8 copies of one of the two split models (pink in Figure 10 (d)). Following MR with refinement in Refmacat and model building using Modelcraft, brought the structure close to completion (R/Rfree=0.25/0.27) but it became clear that the 25 residues were missing from the N-terminus. This absence also meant that the solvent content was higher (67.6%) than was estimated using a Matthews coefficient calculation based on the initial sequence (52%).

**Figure 10.**
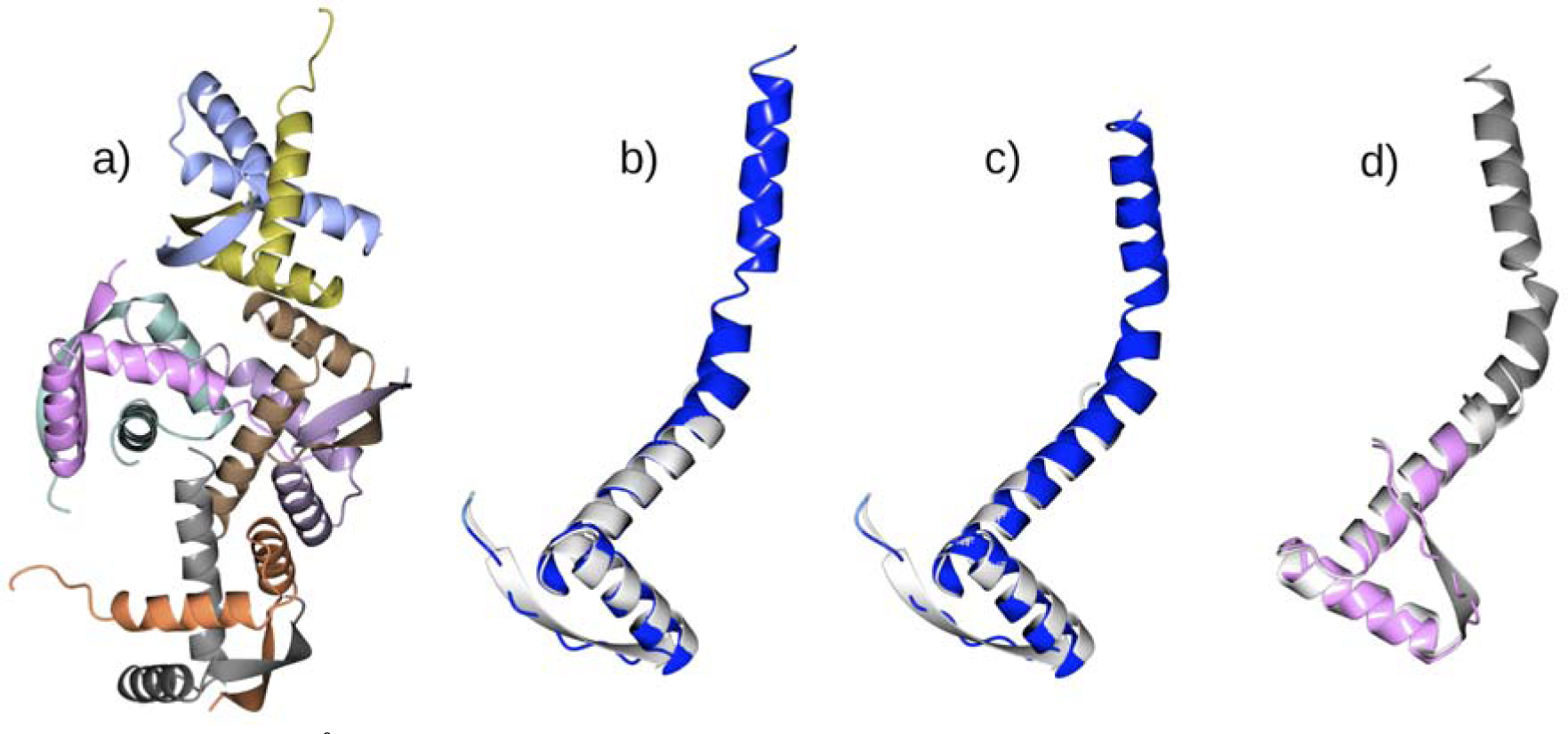
8u12 (1.6Å, 8 copies in the asymmetric unit, spacegroup H3). (a). (b) and (c) show the alignment of the predicted model from AF2 and CF onto a single chain from the target (white) respectively. The predicted models are coloured by pLDDT. Note that the predicted models contain about 25 additional C-terminal residues. These residues from the expected sequence (as deposited in the PDB) did not appear in the crystal. The structure was determined by splitting the model in two and finding 8 copies of the N-terminal domain (d pink).

#### 3.2.7 Some targets resist solution by any approach

After application of the battery of tools described above, some targets were not solved using computational predictions (Table 1; Figure 11). Of course, this does not preclude the possibility of some models being able to solve structures with extensive, expert intervention.

**Figure 11.**
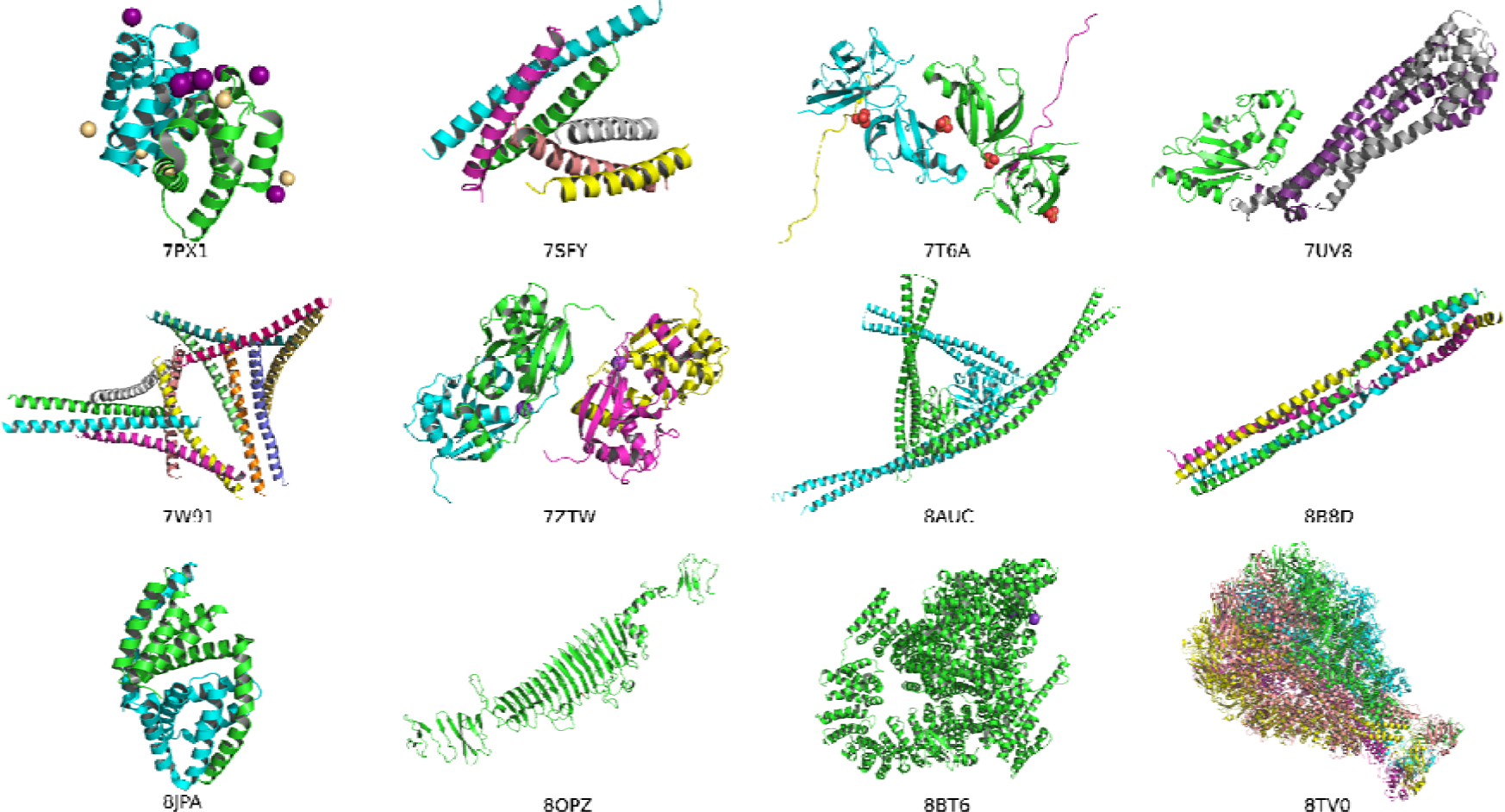
The crystal structures for the 12 cases that we were unable to solve with Colabfold/AlphaFold2 models or alternative MR methods. The structures are coloured by chain.

**Table 1.** The PDB entries we were unable to solve using any MR method.

| PDB ID | Resolution (Å) | Number of molecules in ASU per entity | Biological assembly | Number of residues in each chain | Secondary structure | SOCKET 2: Contains coiled-coil? | AF2 Neff/length | AF2 total sequences in MSA | AF2 Average pLDDT | AF2 TMscore | CF Neff/length | CF total sequences in MSA | CF Average pLDDT | CF TMscore |
| --- | --- | --- | --- | --- | --- | --- | --- | --- | --- | --- | --- | --- | --- | --- |
| **7px1** | 2.33 | 2 | Homo 2-mer | 89 | All helix | No | 0.100 | 19 | 69.2 | 0.630 | 0.030 | 8 | 80.2 | 0.66 |
| **7sfy** | 2.5 | 4, 2 | Hetero 3-mer | 46 | All helix | Yes | 3.99, 1.51 | 375, 190 | 91.2, 87.2 | 0.72, 0.68 | 0.29, 0.67 | 53, 122 | 94.5, 88.4 | 0.72, 0.68 |
| **7t6a** | 1.65 | 2, 2 | Hetero 2-mer | 185, 44 | Mostly sheet, All sheet | No, No | 0.08, 0.58 | 31, 52 | 47, 59.5 | 0.49, 0.33 | 0.02, 0.02 | 13, 2 | 38.4, 71.1 | 0.37, 0.32 |
| **7uv8** | 2.7 | 1, 2 | Hetero 3-mer | 150, 212 | Mostly helix, All helix | No, Yes | 23.23, 2.46 | 12064, 860 | 95.8, 78.4 | 0.98, 0.23 | 61.53, 0.17 | 14569, 64 | 95.2, 79.5 | 0.98, 0.23 |
| **7w91** | 3.29 | 12 | Homo 6-mer | 56 | All helix | Yes | 2.010 | 551 | 94.8 | 0.740 | 1.320 | 655 | 96.1 | 0.72 |
| **7ztw** | 1.93 | 4 | Homo 2-mer | 150 | Mostly helix | No | 7.510 | 1298 | 69.5 | 0.400 | 0.340 | 149 | 78.1 | 0.43 |
| **8auc** | 3.5 | 2 | Monomer | 600 | Mostly helix | Yes | 2.940 | 13886 | 78.9 | 0.340 | 1.170 | 19173 | 76.5 | 0.32 |
| **8b8d** | 2.4 | 4 | Homo 4-mer | 115 | All helix | Yes | 0.160 | 55 | 86.3 | 0.660 | 0.070 | 46 | 83.3 | 0.56 |
| **8jpa** | 2.3 | 2 | Homo 2-mer | 183 | All helix | No | 0.250 | 799 | 85.3 | 0.490 | 0.020 | 91 | 85.8 | 0.48 |
| **8opz** | 1.5 | 1 | Homo 3-mer | 641 | Mostly sheet | No | 0.010 | 349 | 67.3 | 0.640 | 0.010 | 356 | 46.1 | 0.48 |
| **8bt6** | 2.33 | 1 | Monomer | 2646 | Mostly helix | Yes | 0.350 | 13505 | - | - | 0.010 | 103 | 83.3 | 0.73 |
| **8tv0** | 3.1 | 5 | Homo 5-mer | 2537 | Mixed | Yes | 0.280 | 20759 | - | - | 0.150 | 8135 | 79.6 | 0.68 |

Inspection of Table 1 suggests that traditional complications of MR such as moderate diffraction resolution, and the search model comprising only a small proportion of the scattering matter still apply in some cases. So, even a perfect model will fail in cases of many chains in the asymmetric unit (where a relevant biological oligomer can not be accurately built) and insufficient resolution (Oeffner *et al*., 2018).

Here the focus is on model-centred factors that complicate MR. Given the overall excellent performance of AF2 and CF predictions as search models, but the understanding that modelling fails in cases with shallow MSAs, our initial prediction was that this set of recalcitrant targets would comprise largely orphan sequence, or members of small families with few homologues in sequence databases. A rough minimum of 30 sequences in the MSA is often quoted as a threshold for successful modelling (Jumper *et al*., 2021), but the diversity of the MSA is also important, hence the use of Neff/length here, where Neff is the number of effective (non-redundant to 80% sequence identity) sequences and the value is normalised by length. By this measure, 0.2 has been quoted as a threshold below which the MSA may be too shallow for good modelling (Ruiz-Serra *et al*., 2021). The set of failing PDBs in Table 1 covers 12 sequences. Of these, four lie below the Neff/length threshold in AF2 runs and eight in CF runs. For comparison, more than a third of the easily solved set had Neff/length values for CF runs of < 0.2. This observation suggests that shallow MSAs are only part of the explanation of the difficulties encountered with this set.

Interestingly, seven of 12 chains in Table 1 contain coiled-coil regions. This compares to only 42 of 355 (12%) in the set that were easily solved in an automated fashion (Supp Table 1). Furthermore, half of the structures that yielded only to ideal helices or helical ensembles (see above) contained coiled-coil. These observations suggest that, even with the current generation of modelling software, this architecture remains a continuing challenge in MR (Thomas *et al*., 2015; Caballero *et al*., 2018; Thomas *et al*., 2020; Fu *et al*., 2022).

Interestingly, among the most intractable of the recent CASP15 targets (Simpkin, Mesdaghi *et al*., 2023), high α-helix content was also seen as aggravating: indeed six of 14 chains (43%) in the 12 structures in Table 1 are defined as all-alpha. This compares with 42 all-alpha in the set of 356 (12%) that solved readily (Supp Table 1) suggesting that high helical content is indeed linked to AF2 target difficulty.

Finally, it is interesting to note that some targets (7px1, 7t6a, 7ztw) contain multiple metal ions: conceivably these, which are not considered explicitly by AF2, may have a significant influence on conformation as well as comprising, in some cases, a significant proportion of the scattering matter.

As this work was being finalised for publication, AlphaFold 3 (AF3) was released (Abramson *et al*., 2024) and structure solution of the 12 cases in Figure 11 was attempted using AF3 models and the same methods. The only case that readily solved was 8bt6 (Supp Figure 2). AF3 produced a more accurate model that was therefore better split into structural units for MR. Note that in this single case an AF2 model was not available due to system limitations at the modelling stage.

## 4 Conclusions

It is already very apparent that the emergence of AF2 and similar tools has revolutionised the phasing step of X-ray crystallography. Already by far the most favoured approach, MR has been turbocharged by the ever easier availability of high quality structure predictions for most proteins. The literature contains many examples of structures that resisted structure determination until AF2 allowed easy phasing by MR (Barbarin-Bocahu & Graille, 2022).

Larger scale exercises, especially focussed on CASP targets have also shown that most crystal structures can be phased with AF2 models, even those ‘Free Modelling’ targets which have only weak sequence and structural similarities with PDB entries (Millán *et al*., 2021; Pereira *et al*., 2021; McCoy *et al*., 2022; Simpkin, Mesdaghi *et al*., 2023).

In this work we focused on structures that were determined by SAD in the period July 2022 to October 2023. It is important to note that these are not representative of the PDB as a whole: in the same period, around 88% of structures were determined by MR. While there may be other reasons to choose SAD phasing e.g. determination of bound ion positions, our focus is on a set of targets that is likely to include most targets that were not easily tractable by MR at the time. In this way, we hoped to explore outer boundaries of phasing with AF2 and other bioinformatics methods, and to reveal how many structures and what kind of structure require experimental phasing methods post-AF2. Notably, AF3, which became available at a very late stage, does not seem to change the picture dramatically with regard to providing search models for MR.

Our results show that a striking 356/406 or 87.7% of the set could be solved directly using an AF2 model, unprocessed except for conversion of pLDDT confidence estimates to B-factors. In a further number of cases, elementary model editing to remove low-confidence regions, a procedure available in major software packages, enables structure solution. Also now routine, splitting of a structure prediction into individual domains (Oeffner *et al*., 2022; Simpkin *et al*., 2022) was sometimes required for structure solution: however, in some cases success depended on the particular splitting algorithm used. In other cases, splitting domains was not essential but provided solutions that were significantly better quality, accelerating downstream rebuilding and refinement.

Use of methods other than AF2 was sometimes important. In four cases, we found that neither AF2 flavour succeeded, but an edited model from the protein Language Model method ESMFold (Lin *et al*., 2023) succeeded. In three other cases, construction of a multimeric search model was required for success. As with search model splitting, the use of a multimeric search model was sometimes advantageous for speed and for the better phases that resulted, even when a monomeric search model could solve a structure.

Interestingly, oligomeric state can be predicted to some extent using Deep Learning networks (Kshirsagar *et al*., 2024). In four further cases, tertiary structure prediction failed to produce a successful search model, but MR with secondary structure elements, using ARCIMBOLDO (Rodríguez *et al*., 2009) or AMPLE (Sánchez Rodríguez *et al*., 2020), succeeded, emphasising the continued utility of these methods in some cases.

Ultimately, these approaches collectively failed on 12 targets. One relevant feature is low MSA depth i.e. some of these proteins have few homologues in the databases. This feature deprives AF2 of the evolutionary covariance information which (in the absence of templates, of course) is essential for construction of an initial model (Jumper *et al*., 2021). This mirrors other findings, such as those of recent CASPs (Pereira *et al*., 2021; Simpkin, Mesdaghi *et al*., 2023). This remains a current development frontier of predictive methods despite the possibilities of artificial MSA depth enhancement (Zhang *et al*., 2023) and means to recognise the fast evolution of viral proteins and detect more distant homologues than current pipelines do (Karlin, 2024). There are also recurrent suggestions that pLM-based methods may outperform MSA-based approaches in such cases (Chowdhury *et al*., 2022; Wang *et al*., 2022; Jing *et al*., 2024). although this was not apparent at the CASP15 experiment (Simpkin, Mesdaghi *et al*., 2023).

Also notable are the continued difficulties with coiled-coil proteins, whose unpredictable irregularities, oligomeric state and parallel vs antiparallel architectures remain, despite ongoing efforts (Madeo *et al*., 2023; Madaj *et al*., 2024), a significant hindrance to structure prediction and hence the use of modelled structures for MR.

Of course, a sufficiently accurate search model is necessary but not sufficient for structure solution by MR. Even with a good model available, a case may still fail where one or more traditional complications apply. These include cases where a large number of copies of the search model are present in the asymmetric unit and crystals that diffract only to poorer resolution. Thus, in terms of prioritising cases for experimental phasing beamlines, difficult targets can initially be predicted bioinformatically, especially by assessing the depth and diversity of the MSA that can be constructed from homologous sequences, but other complications will only become evident once crystal characteristics can be seen.

