## Supplementary Material for "In the AlphaFold era, when is experimental phasing of protein crystals still required?"

When is experimental phasing of protein crystals needed for structure solution, post-AlphaFold 2?

### PDB IDS:

6st5, 7ej3, 7el5, 7eoz, 7ewf, 7fa1, 7fac, 7fax, 7fex, 7fgn, 7fgp, 7fh5, 7fia, 7fib, 7mch, 7o5i, 7o6t, 7o6u, 7o6v, 7o6w, 7p4a, 7pc1, 7pcv, 7pjr, 7pk2, 7pv5, 7pv7, 7pva, 7pwe, 7px1, 7pzt, 7q5y, 7qce, 7qda, 7qe1, 7qeh, 7qld, 7qlr, 7qrj, 7qrr, 7qsj, 7qwa, 7qwb, 7qwc, 7qwd, 7qwe, 7qx4, 7r0t, 7r0x, 7r1m, 7r6s, 7rdn, 7ris, 7rkb, 7rl3, 7roa, 7rt7, 7rxu, 7sfn, 7sfy, 7sie, 7smg, 7snr, 7sua, 7sz2, 7sz3, 7t5t, 7t69, 7t6a, 7tdr, 7tds, 7te2, 7th0, 7tlv, 7tlw, 7tom, 7tvl, 7twd, 7u01, 7u08, 7u2v, 7uav, 7uba, 7udi, 7uf8, 7ug8, 7ujb, 7ulo, 7ups, 7ur2, 7urp, 7usn, 7uv8, 7uvg, 7uwu, 7uyg, 7uzt, 7v0f, 7v1o, 7v2s, 7v3o, 7v52, 7v54, 7v56, 7v57, 7v7x, 7vbo, 7vch, 7vep, 7viv, 7vqk, 7vs2, 7vs6, 7vwv, 7vxt, 7vyu, 7w2o, 7w2q, 7w54, 7w63, 7w6y, 7w6z, 7w86, 7w91, 7wa9, 7wdt, 7wej, 7wh9, 7wjg, 7wjl, 7wlh, 7wmw, 7wmx, 7wmy, 7wmz, 7wn7, 7wrw, 7wuk, 7wup, 7wuw, 7wux, 7wuz, 7ww2, 7ww4, 7wwq, 7wwt, 7wwy, 7wx0, 7wx1, 7wzv, 7x0i, 7x0j, 7x0k, 7x0l, 7x0m, 7x0n, 7x0o, 7x15, 7x45, 7x6z, 7x7i, 7x8u, 7x9r, 7xbj, 7xcc, 7xds, 7xfp, 7xg9, 7xgt, 7xhz, 7xky, 7xmw, 7xn2, 7xp9, 7xpc, 7xpi, 7xpr, 7xrb, 7xre, 7xrx, 7xzf, 7y0y, 7y19, 7y3w, 7y56, 7y78, 7y79, 7y7o, 7y8u, 7yco, 7ydo, 7yfj, 7ygf, 7yhl, 7yik, 7yji, 7yjp, 7yjq, 7yjr, 7yjs, 7yjt, 7yk4, 7ykv, 7yle, 7ym5, 7ym7, 7ymo, 7yn1, 7ynx, 7ype, 7ypf, 7yr9, 7yt9, 7ytl, 7ytt, 7ytu, 7yuj, 7yv0, 7z3b, 7zbh, 7zgi, 7zhd, 7zhl, 7zju, 7zk1, 7znr, 7zns, 7ztw, 7zu3, 7zu8, 7zug, 7zv1, 7zyh, 8a14, 8a1i, 8a24, 8a2n, 8a30, 8a38, 8a82, 8aaj, 8ab2, 8abt, 8aew, 8aez, 8af9, 8ag9, 8ahd, 8ahz, 8aid, 8ajq, 8am4, 8asa, 8au6, 8auc, 8avz, 8ax2, 8ay2, 8b2e, 8b3e, 8b3w, 8b4l, 8b55, 8b73, 8b8d, 8bc5, 8bcx, 8bd1, 8bfh, 8bfi, 8bgt, 8bhd, 8bjw, 8bkd, 8bke, 8brp, 8bt6, 8bve, 8bvl, 8bvp, 8c3d, 8car, 8cgm, 8cnr, 8cpn, 8cwt, 8d0o, 8d3t, 8d89, 8da2, 8dc1, 8deh, 8df2, 8dfk, 8dop, 8dp6, 8dpk, 8dq2, 8dtn, 8dtq, 8dtu, 8dvq, 8dwz, 8ebf, 8ebg, 8efm, 8ehc, 8em5, 8emb, 8en9, 8ena, 8eo2, 8ep6, 8ewh, 8ezo, 8ezp, 8ezr, 8ezs, 8ezu, 8ezx, 8f00, 8f01, 8f03, 8f05, 8f06, 8f07, 8f0b, 8f3k, 8f7n, 8f8n, 8fbe, 8fby, 8fia, 8fnr, 8fns, 8fzz, 8g1n, 8g1y, 8g28, 8gbe, 8gj9, 8gjw, 8gjy, 8glb, 8gq9, 8gs1, 8gsx, 8gt9, 8gup, 8gxl, 8gy4, 8gy8, 8h0h, 8h0r, 8h1e, 8h1f, 8h1g, 8h3z, 8h8h, 8hav, 8haw, 8hbr, 8hd2, 8hdv, 8heh, 8hek, 8hhv, 8hja, 8hn2, 8hn3, 8hp8, 8hx3, 8i16, 8i2d, 8i3j, 8i59, 8i6h, 8i6z, 8i8y, 8ic1, 8iib, 8ilc, 8j1w, 8j1x, 8j67, 8j8p, 8j8q, 8j98, 8jj7, 8jpa, 8k1c, 8k1f, 8k1i, 8k5l, 8k76, 8ok4, 8opz, 8oq1, 8pfc, 8siu, 8smq, 8srz, 8tv0, 8u00, 8u01, 8u12

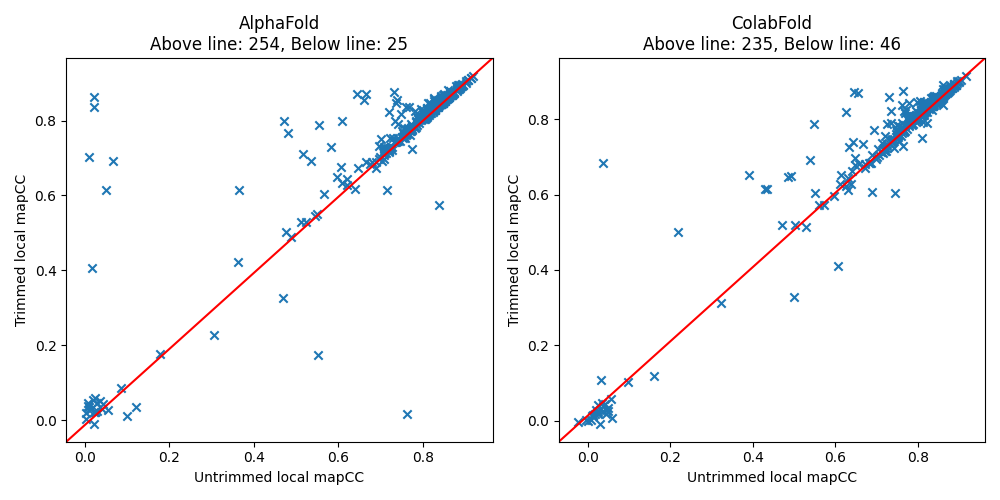

Supplementary Fig 1. A comparison of the local mapCC for AlphaFold2 and ColabFold models before and after trimming residues with a pLDDT <70. Points above the red line indicate that trimming has improved the mapCC and points below the red line indicate that trimming has decreased the mapCC.

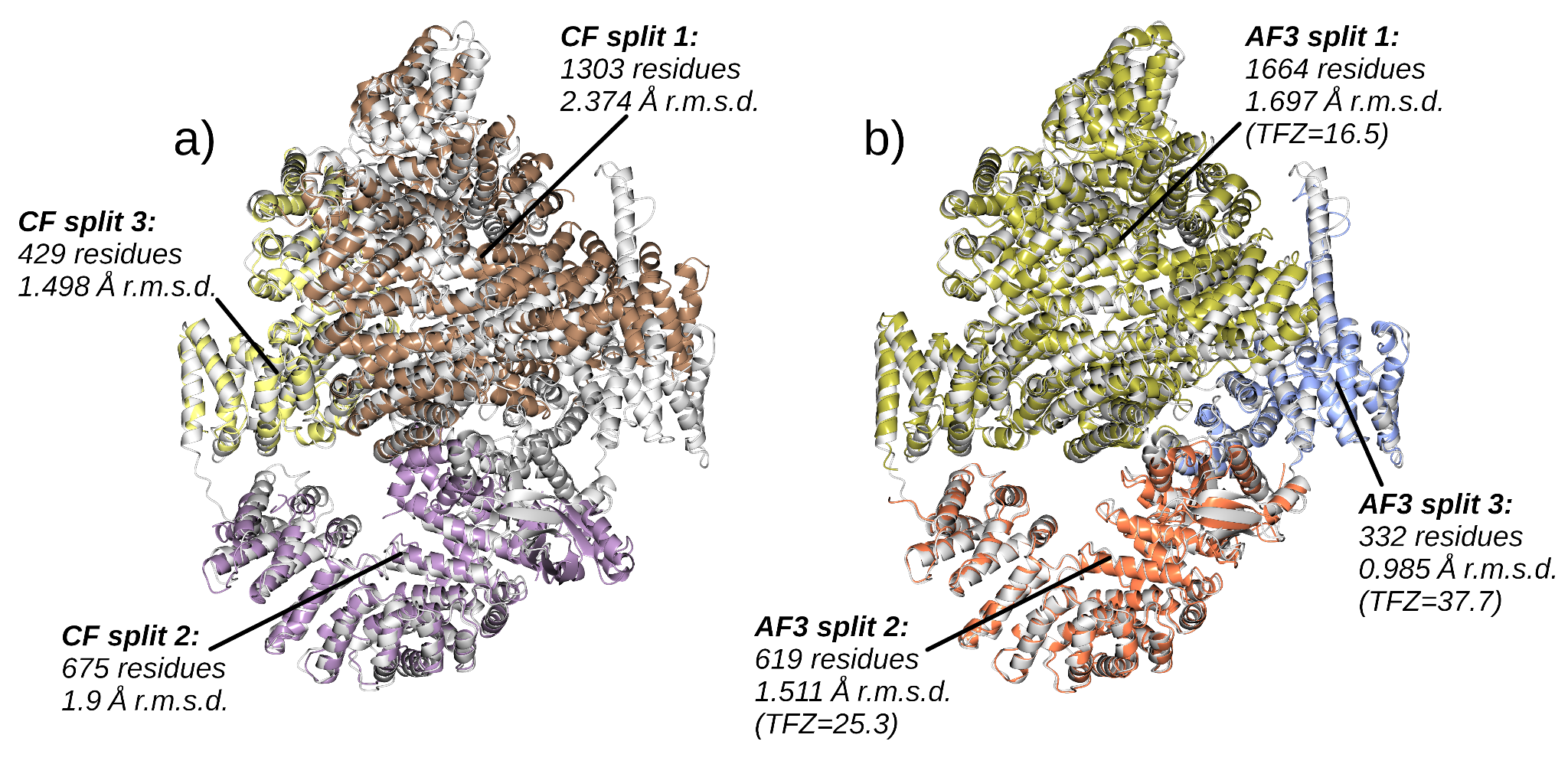

Supplementary Fig 2 8bt6 (2.33Å, 1 copy in the asymmetric unit, spacegroup P212121). The CF (a) and AF3 (b) predictions split 3-ways by SnD and aligned to target (white) by Gesamt. Number of residues and r.m.s.d. to the target for each part are shown. Phaser was not successful in attempting to place the 3 search models created from the CF prediction but was able to place all 3 search models from the more accurate AF3 prediction with an LLG of 1794 (TFZ scores for each model are shown). Subsequent refinement of the AF3 solution resulted in an R/Rfree of 0.25/0.32. Due to the size of the structure, we were unable to predict the target structure using AF2 as we lacked access to the computational resources needed. The AF3 prediction was generated using the Alphafold server (Abramson *et al.*, 2024).

[Abramson, J., Adler, J., Dunger, J., Evans, R., Green, T., Pritzel, A., Ronneberger, O., Willmore, L., Ballard, A. J., Bambrick, J., Bodenstein, S. W., Evans, D. A., Hung, C.-C., O’Neill, M., Reiman, D., Tunyasuvunakool, K., Wu, Z., Žemgulytė, A., Arvaniti, E., Beattie, C., Bertolli, O., Bridgland, A., Cherepanov, A., Congreve, M., Cowen-Rivers, A. I., Cowie, A., Figurnov, M., Fuchs, F. B., Gladman, H., Jain, R., Khan, Y. A., Low, C. M. R., Perlin, K., Potapenko, A., Savy, P., Singh, S., Stecula, A., Thillaisundaram, A., Tong, C., Yakneen, S., Zhong, E. D., Zielinski, M., Žídek, A., Bapst, V., Kohli, P., Jaderberg, M., Hassabis, D. & Jumper, J. M. (2024). *Nature* **630**, 493–500.](http://paperpile.com/b/mpGRWR/ICjV)
